# Studies on the mechanism of membrane mediated general anesthesia

**DOI:** 10.1101/313973

**Authors:** Mahmud Arif Pavel, E. Nicholas Petersen, Hao Wang, Richard A. Lerner, Scott B. Hansen

## Abstract

Inhaled anesthetics are a chemically diverse collection of hydrophobic molecules that robustly activate TWIK related K+ channels (TREK-1) and reversibly induce loss of consciousness. For a hundred years anesthetics were speculated to target cellular membranes, yet no plausible mechanism emerged to explain a membrane effect on ion channels. Here we show that inhaled anesthetics (chloroform and isoflurane) activate TREK-1 through disruption of palmitate-mediated localization of phospholipase D2 (PLD2) to lipid rafts and subsequent production of signaling lipid phosphatidic acid (PA). Catalytically dead PLD2 robustly blocks anesthetic TREK-1 currents in cell patch-clamp. Localization of PLD2 renders the anesthetic-insensitive TRAAK channel sensitive. General anesthetics chloroform, isoflurane, diethyl ether, xenon, and propofol disrupt lipid rafts and activate PLD2. In the whole brain of flies, anesthesia disrupts rafts and PLD^null^ flies resist anesthesia. Our results establish a membrane mediated target of inhaled anesthesia and suggest PA helps set anesthetic sensitivity *in vivo*.

## INTRODUCTION

In 1846 William Morton demonstrated general anesthesia with inhaled anesthetic diethyl ether (*1*). For many anesthetics (but not all), lipophilicity is the single most significant indicator of potency; this observation is known as the Meyer-Overton correlation (*2, 3*). This correlation, named for its discoverers in the late 1800’s, and the chemical diversity of anesthetics (Supplementary Fig. S1a) drove anesthetic research to focus on perturbations to membranes as a primary mediator of inhaled anesthesia (*3*). Over the last two decades, enantiomer selectivity of anesthetics suggested a chiral target and direct binding to ion channels emerged (*4*). But the possibility of a membrane mediated effect has remained (*5–7*).

In 1997 a theory emerged suggesting that disruption of ordered lipids surrounding a channel could activate the channel (*8*). Disruption of ordered lipids (sometimes referred to as lipid rafts (*9*)) allows phospholipase D2 (PLD2) to translocate out of lipid domains and experience a new chemical environment (*10*) (Supplementary Fig. S1b-c). PLD2 actives Twik related potassium channel-1 (TREK-1). If inhaled anesthetics can disrupt lipid domains to activate a channel, this would constitute a mechanism distinct from the usual receptor-ligand interaction and establish a definitive membrane mediated mechanism for an anesthetic.

TREK-1 is an anesthetic-sensitive two-pore-domain potassium (K2P) channel. Xenon, diethyl ether, halothane, and chloroform robustly activate TREK-1 at concentrations relevant to their clinical use (*11, 12*) and genetic deletion of TREK-1 decreases anesthesia sensitivity in mice (*13*). Here we show inhaled anesthetics disrupt palmitate-mediated localization in the membranes of cultured neuronal, muscle cells, and in the whole brain of anesthetized flies. The disruption releases PLD2 from muscle and neuronal cells which signal downstream to activate TREK-1 channels. This result establishes the membrane as a pertinent target of inhaled anesthetics.

### Anesthetics perturb nanoscale lipid heterogeneity (GM1 domains)

The best studied lipid domains contain saturated lipids cholesterol and sphingomyelin (e.g. monosialotetrahexosylganglioside1 (GM1)) (see Supplementary Fig. S1b-c) and bind cholera toxin B (CtxB) with high affinity (*9*). Anesthetics lower the melting temperature and expand GM-1 domains in artificial membranes and membrane vesicles (*14–16*). Lipids are thought to exist in heterogenous mixtures with features smaller than 100 nm that are best characterized with super-resolution imaging (e.g., direct stochastical optical reconstruction microscopy (dSTORM) and stimulated emission depletion (STED)) (*17–20*). Palmitoylation, a posttranslational modification that covalently attaches a 16 carbon lipid, localizes proteins to GM1 lipids (*21*) (Supplementary Fig. S1c) including many ion channels (*22*). We used dSTORM to test the hypothesis that anesthetics harness lipid heterogeneity to gate TREK-1 (*23*).

To test an anesthetic specific perturbation to GM1 lipids, we treated neuroblastoma 2A (N2A) cells with anesthetic chloroform at 1mM concentration and monitored fluorescent CtxB clustering by dSTORM (Fig. 1a). Chloroform strongly increased both the diameter and area of GM1 domains in the cell membrane (Fig. 1b-c, Supplementary Fig. S2a). Isoflurane (1 mM) behaved similar chloroform (Fig. 1a-c). The Ripley’s radius, a measure of space between domains, decreased dramatically for both chloroform and isoflurane (Supplementary Fig. S2b) suggesting the domains expand (*24*) and possibly divide as the number of rafts per cell increased (Fig. 1d-e, Supplementary Fig. S2c). Methyl-β-cyclodextrin (MβCD), a chemical that disrupts GM1 domains by removing cholesterol (*10*), reduced the total number of domains (Fig.1d). Binning the domains into small (0-150 nm) and large (150-500nm) revealed a clear shift from small to large domains in the presence of inhaled anesthetics and revealed the opposite effect after MβCD treatment (Supplementary Fig. S2d). Similar results were obtained in C2C12 myoblasts (muscle cells); chloroform strongly increased both the diameter and area of GM1 domains. (Supplementary Fig. S2e-h)

**Figure 1.**
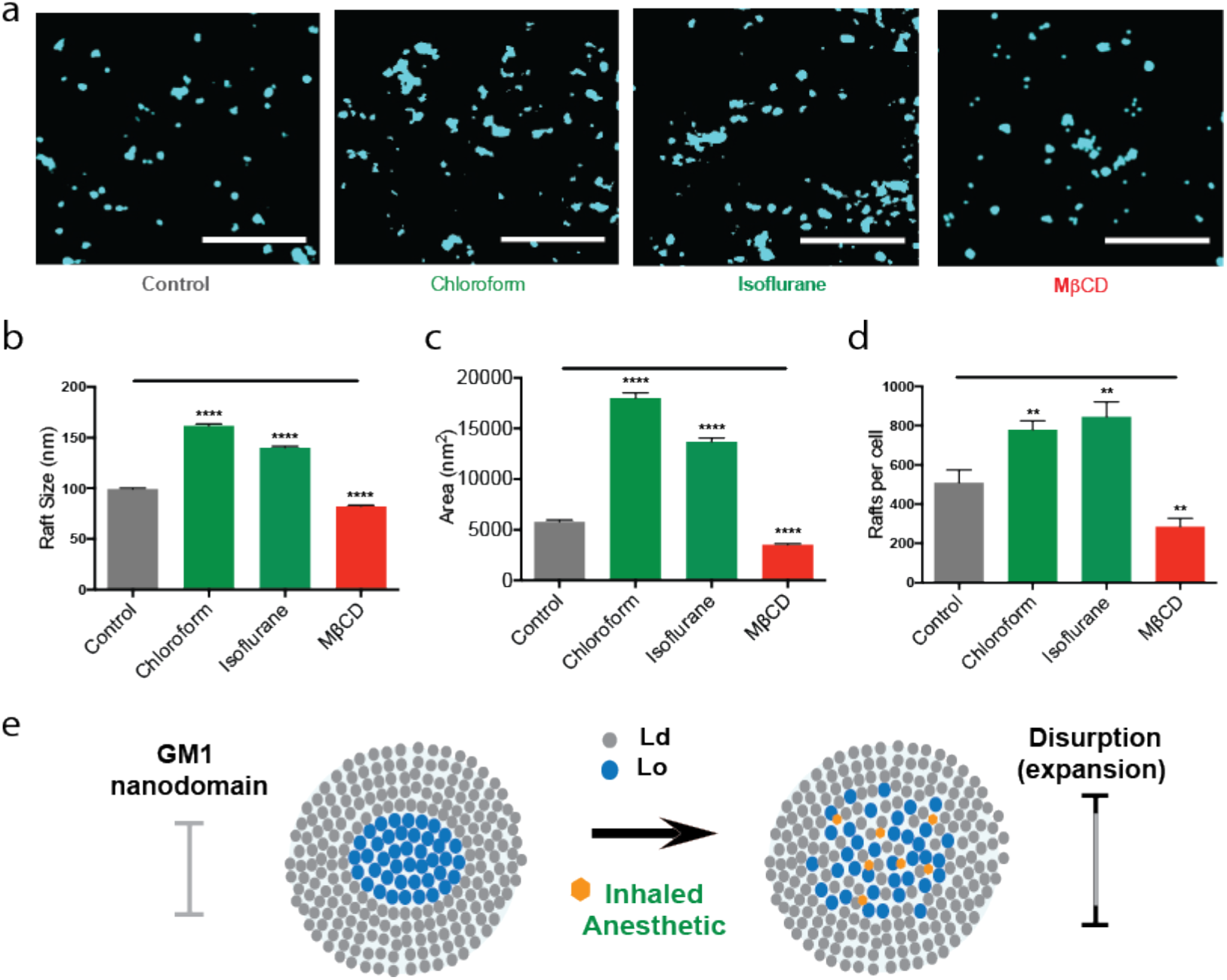
Inhaled anesthetics disrupt GM1 domain structure. **a**, Representative reconstructed super-resolution (dSTORM) images of GM1 domains (lipid rafts) before and after treatment with chloroform (1 mM), isoflurane (1 mM), or MβCD (100 µM) (Scale bars: 1 µm). **b-d**, Bar graphs comparing the average sizes (**b**) and areas (**c**) quantified by cluster analysis (± s.e.m., n = 2842-7382). **d**, Quantified number of rafts per cell. (± s.e.m., n=10) (Student’s t-test results: ****P<0.0001) **e**, Model representation of raft disruption by anesthetics. GM1 lipids (blue) form ordered domains. Inhaled anesthetic (orange hexagon) intercalate and disrupt lipid order causing the domain to expand.

### Mechanism of anesthetic sensitivity in TREK-1 channels

To confirm the observed disruption is capable of producing a biological signal, we studied TREK-1 activation by inhaled anesthetics. Activation of TREK-1 by inhaled anesthetics, was previously shown to require a disordered loop in the channel’s C-terminus (*11*) (Supplementary Fig. S3a-b). The enzyme phospholipase D2 (PLD2) also binds to and activates through the same C-terminal region in TREK-1 (*25*). PLD2 is palmitoylated at cysteines near its Pleckstrin homology (PH) domain, which is required to localize it to GM1 lipids (*26*). The PH domain also binds PIP_2_ which opposes the localization by palmitoylation (Supplementary Fig. S1c). Disruption of GM1 lipids by mechanical force activates PLD2 by substrate presentation—the enzyme translocated from GM1 to PIP_2_ lipids near its substrate (*10*). If anesthetics disrupt GM1 domains, then we expect PLD2 or a complex of PLD2 and TREK-1 to translocate and activate TREK-1. If correct, the PLD2 dependent TREK-1 activation would confirm lipid disruption as a component of TREK-1 anesthetic sensitivity and a membrane mediated pathway.

To test our hypothesis and the role of lipid heterogeneity in anesthetic sensitivity of TREK-1, we applied chloroform to HEK293 cells over-expressing TREK-1 and a catalytically dead K758R PLD2 mutant (xPLD2) that blocks anionic lipid production (e.g. PA and PG) (*27*). We over expressed TREK-1 in HEK293 cells to avoid the confounding effects of endogenous channels other than TREK-1 found in N2A and C2C12 cells. We found xPLD2 blocked all detectible chloroform specific current (Fig. 2a-c). Cells not transfected with TREK-1 had no observable current and were unaffected by chloroform treatment (Supplementary Fig. S3c) showing the xPLD’s effect is specific to TREK-1 currents. We confirmed xPLD overexpression with western blot analysis (Supplementary Fig. S3d). This result suggests a two-step mechanism for anesthetic action on TREK-1 channels. First, anesthetics disrupt GM1 domains releasing PLD2 and second, the enzyme binds to the C-terminus of TREK-1 and they translocates to substrate and activates the channel through increased local concentration of anionic lipid (Fig. 2d). Similarly, PLD could be pre-bound to TREK-1 and translocation as a complex to PIP_2_ domains and PLD substrate.

**Figure 2.**
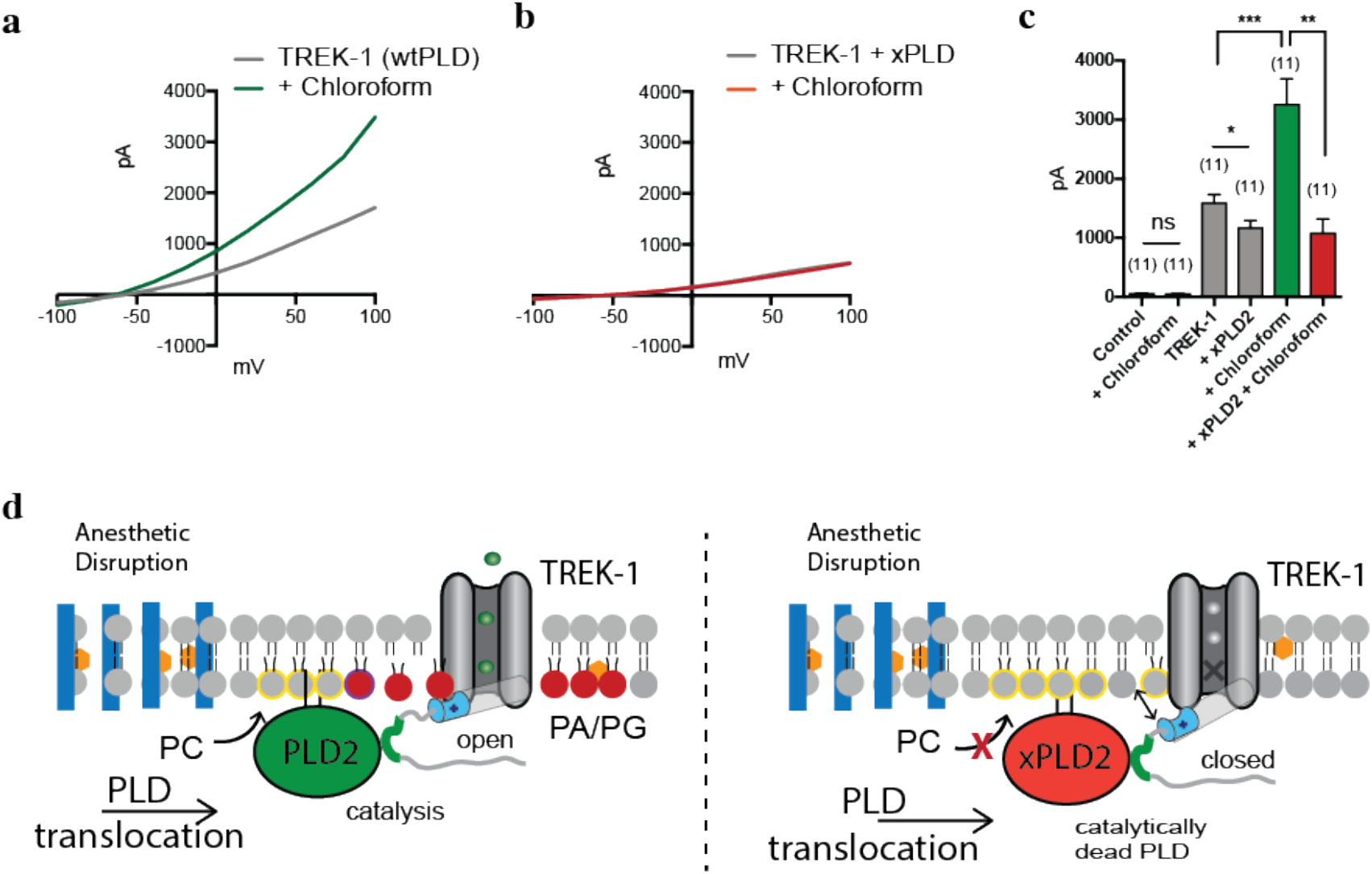
Activation of TREK-1 by inhaled anesthetic is PLD2 dependent. **a**, Representative TREK-1 whole-cell currents activated by chloroform (1 mM) in physiological K^+^ gradients. The current-voltage relationships (I-V curves) were elicited by 1-s depolarizing pulses from −100 to 100 mV in +20 mV increments. **b**, Representative I-V curves showing that co-expression of a catalytically inactive mutant of PLD2 (xPLD2 = PLD2_K758R) abolishes the TREK-1 activation by chloroform. **c**, Bar graph showing the ∼2-fold increase of TREK1 current when activated by chloroform (1 mM) (n = 11) at +40 mV (± s.e.m.). **d,** Schematic representation of TREK-1 activation by inhaled anesthetics. Anesthetic disruption of GM1 domains causes PLD2 to localize with TREK-1 and its substrate phosphatidylcholine (PC) in the disordered region of the membrane. As PLD2 hydrolyzes PC to phosphatidic acid (PA), the anionic lipid binds to a known gating helix (grey cylinder), with a lipid binding site (cyan)(*30*), that activates TREK-1. Student’s t-test results: *P < 0.05; **P<0.01; ***P<0.001; NS ≥ P.0.05.

### Transfer of anesthetic sensitivity to TRAAK

TWIK-related arachidonic acid-stimulated K+ channel (TRAAK) is an anesthetic insensitive homolog of TREK-1 (Supplementary Fig. S3e). Interestingly, native TRAAK is also insensitive to PLD2 (*25*). However, concatenating PLD2 to the N-terminus maximally activates TRAAK and introduction of the PLD2 binding domain from TREK-1 renders TRAAK PLD2 sensitive (*25*). If PLD2 is responsible for anesthetic sensitivity in TREK-1, we reasoned we could render TRAAK anesthetic sensitive by introducing the PLD2 binding site into the C-terminus of TRAAK (Fig. 3a).

**Figure 3.**
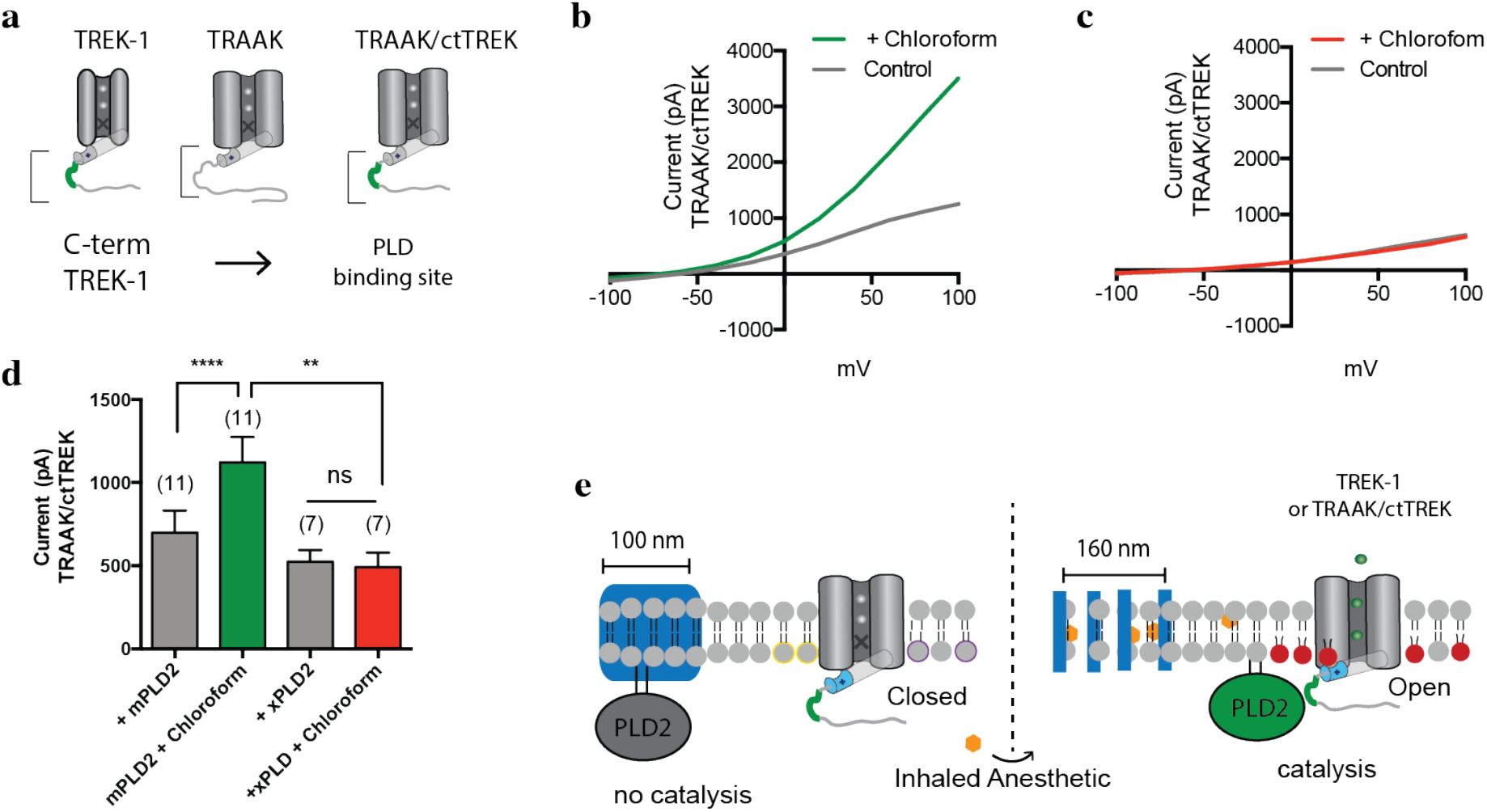
PLD2 localization renders TRAAK anesthetic sensitive. Native TRAAK is an anesthetic insensitive channel. **a,** Cartoon showing the experimental setup. TRAAK is fused with the C-terminus of TREK-1 (TRAAK/ctTREK). The BLD binding site is depicted in green. **b-c**, Representative I-V curve showing TRAAK/ctTREK-1 is activated by chloroform when co-expressed with mouse PLD2 (mPLD2) (**b**). The co-expression of the catalytically inactive PLD2 (xPLD2) abolishes the chloroform activation of TRAAK/ctTREK-1 chimeric channel (± s.e.m., n = 7) (**c**). **d,** Bar graph summarizing TRAAK/ctTREK-1 chimeric channel current in the presence or absence of xPLD2 and chloroform (1mM) at +40 mV (± s.e.m., n = 11) (Student’s t-test results: ****P<0.0001, **P<0.01; NS ≥ P.0.05.) **e,** Model mechanism showing that anesthetics activate the TRAAKctTREK-1 chimeric channel through raft disruption and PLD2 substrate presentation; xPLD2 abolishes the activation (the color scheme is as in Fig. 2).

We over expressed the previously characterized PLD2 sensitive TRAAK chimera (*25*) (TRAAK/ctTREK) in HEK cells. As expected, in the presence of 1mM chloroform, TRAAK/ctTREK robustly responded to chloroform (Fig. 3b-d). To confirm the response is due to PLD2 localization and not a direct interaction of the anesthetic with a structural feature of the TREK-1 C-terminus, we over expressed the chimera with xPLD2 and found chloroform had no effect on the channel. This result suggests the disordered C-terminus exerts its anesthetic effect through binding to PLD2 and not direct binding of anesthetic to the C-terminus (Fig. 3e).

The lack of TREK-1 current in the presence of anesthetic (Fig 2b-c) suggests direct binding of anesthetic is insufficient to activate the channel absent PLD2 activity. This result opposes what was previously thought, that the anesthetic directly activates TREK-1(*28*). To confirm our result and distinguish the contribution of an indirect anesthetic effect from a direct one, we tested the contribution of direct binding to TREK-1 anesthetic sensitivity in a flux assay. Ion channels are routinely expressed, purified, and assayed in vesicles and purified TREK-1 channels robustly recapitulates small molecules binding to channels using a flux assay in a cell free system (*29–31*).

Functionally reconstituted TREK-1 in 16:1 phosphatidylcholine (PC) liposomes with 18:1 phosphatidylglycerol (PG) (85:15 mol% ratio) was unaffected by chloroform or isoflurane (1 mM) (Supplementary Fig. S3f-g). Changing the lipids and ratios to 18:1 PC/18:1 PG (90/10 mol%)(*30*) also had no effect (data not shown). To assure that the channel was properly reconstituted and in conditions capable of increased potassium flux, we reconstituted a mutant TREK-1 with double cysteines that covalently lock TREK-1 in the activated state (*30, 32*). Compared to the open TREK-1 control, inhaled anesthetics failed to activate TREK-1 (Supplementary Fig. S3f-g). We showed in a previous study, using the same assay, a clear direct inhibition with local anesthetics (*31*).

### Anesthetics displace PLD2 out of GM1 lipids

For raft disruption to be biologically significant it must transduce the disruption into a biological signal. To test our hypothesis that the disruption gives rise to PLD2 activity and TREK-1 activation, we directly imaged palmitate-mediated PLD2 localization using dSTORM imaging and anesthetic induced translocation of PLD2 out of GM1 lipids. We measure translocation, by calculating a pairwise correlation of PLD2 with CTxB before and after anesthetic treatment. Pairwise correlation avoids potential artifacts from over sampling, and the limits of the user selected parameters used when determining raft size (Supplementary Fig. S4). Both N2A and C2C12 endogenously express TREK-1 and PLD2 (see methods) allowing us to characterize their translocation under endogenous promoters. TREK-1 is not palmitoylated, but palmitoylation could sequester PLD2 away from TREK-1 (Supplementary Fig. S1b-c) or sequester the pre-bound PLD2/TREK-1 complex into GM1 domains away from PLD’s activating lipid (PC) or both (*10*).

Treating the N2A cells with chloroform or isoflurane (1 mM), caused PLD2 to translocate away from GM1 lipids (Fig 4a). Pair correlation analysis showed PLD2 strongly associated with GM1 domains prior to anesthetic treatment (Fig 4b, grey trace) but only weakly associated after treatment (green traces). This highly significant anesthetic-induced change in PLD2 localization was true across all sizes of GM1 domains ((Fig. 4b-c), pair correlation analysis) suggesting the energetics of PLD2 partitioning into GM1 lipids changed, not merely a change in the size of the domain. The anesthetic-specific translocation was similar in magnitude to MβCD stimulated translocation (Fig. 2b-c) and implies that anesthetic expansion of GM-1 domains has the same effect on palmitate-mediated localization. Expansion is therefore a form of raft disruption (Fig. 4d). We obtained similar pair correlation results in C2C12 cells treated with chloroform (Supplementary Fig. S5). The analysis of pairwise correlation with and without anesthetic, that are otherwise treated identical, also largely rules out artifacts from antibody or toxin labeling or permeabilization as the differences in PLD2 localization.

**Figure 4.**
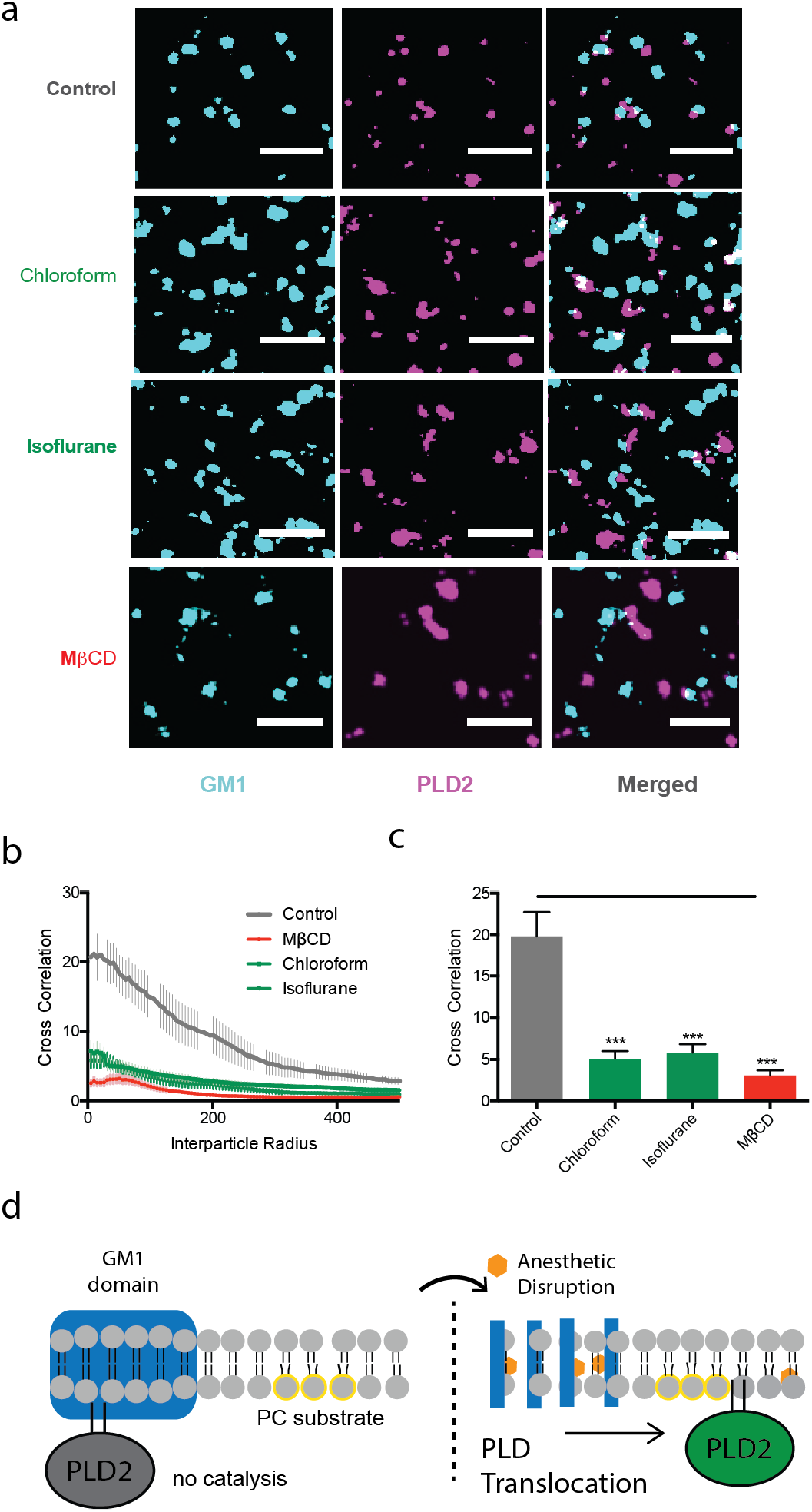
Inhaled anesthetics displace PLD2 from GM1 domains. **a**, Representative super-resolution (dSTORM) images of fluorescently labeled CTxB (lipid raft, from Fig. 1a) and PLD2 before treatment (Control) and after treatment with chloroform (1 mM), isoflurane (1mM), and MβCD (100 µM) in N2A cells (scale bars: 1 µm). **b**, Average cross-correlation functions (C(r)) showing a decrease in PLD2 association with ordered GM1 domains after treatment with anesthetic or MβCD. **c**, Comparison of the first data point in (**b**) (5 nm radius) (± s.e.m., n = 10-17). **d**, Schematic representation of PLD2 in GM1 domain before (left) and after (right) anesthetic treatment. Palmitoylation drives PLD into GM1 domains (blue) away from its unsaturated PC substrate (yellow outline). Anesthetics (orange hexagon) disrupts GM-1 domains causing the enzyme to translocate where it finds its substrate PC in the disordered region of the cell.

### Anesthetics activate PLD2 through raft disruption

If the membrane is a general mechanism for TREK-1 mediated anesthesia, then most known activators of TREK-1 should also activate PLD2. We tested enzymatic activation of PLD2 by treating live cells with a spectrum of chemically diverse inhaled anesthetics and monitoring activity using an assay that couples PLD’s choline release to a fluorescent signal (*10*) (Fig. 5a-e). Diethyl ether, chloroform, isoflurane, and xenon all significantly activated PLD2 in N2A (Fig. 5a-e,h) and C2C12 cells (Supplementary Fig. S6a-d,f). Isoflurane had the greatest effect (Fig. 5h) in N2A cells and chloroform had the greatest effect among inhaled anesthetics in C2C12 cells (Supplementary Fig. S6b,f). The dose response of chloroform and isoflurane reveals an EC50 of 1.0 ± 0.56 mM and 0.8 ± mM respectively (Fig. 5i) in N2A cells. The Hill slopes were 2.4 ± 2.8 and 1.3 ± 0.89 respectively. Ketamine, an injectable NMDA receptor specific anesthetic (*33*) and F6, a non-immobilizer that defies the Meyer-Overton rule, had no effect on PLD activity, as expected (Fig. 5f-h). Dose responses at low concentration were highly variable making the determination of the Hill slope difficult to determine. The PLD2 assay also lacks the ability to mimicked partitioning into the membrane from continuous inhalation.

**Figure 5.**
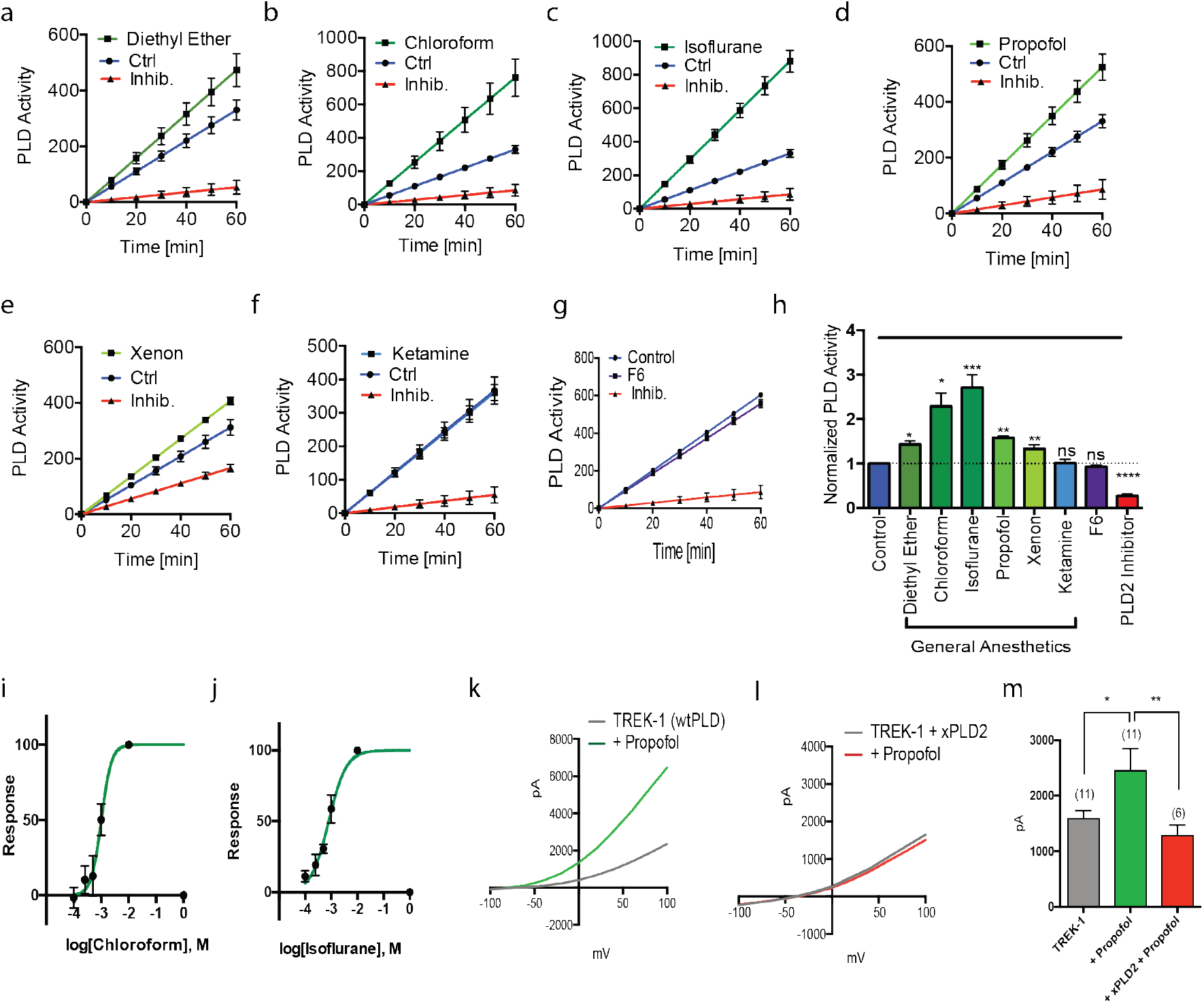
General anesthetics activates phospholipase D2 (PLD2) through raft disruption. (**a-e**) Live cell assays showing the effect of anesthetics on PLD2 activity in N2A cells. Chloroform (1 mM) (**a**), isoflurane (1 mM) (**b**), diethyl ether (1 mM) (**c**), propofol (50 µM) (**d**) and xenon (0.044 µM) (**e**) increased the PLD2 activity as compared with the control cells. Ketamine (50 µM) (**f**) and the non-immobilizer F6 (**g**) had no effect on the PLD2 activity and the activity was inhibited by a PLD2 specific inhibitor (2.5-5 µM) (mean ± s.e.m., n = 4). **h**, Summary of normalized anesthetic induced activity of PLD2 in (a-g) at 60 min. (mean ± s.e.m., n = 4). **i-j**, Dose response of chloroform (EC_50_ = ∼1.0 mM) (**i**) and isoflurane (EC_50_ = ∼0.8 mM) (**j**) on PLD2 activity through raft disruption. **k-l,** Representative I-V curves showing the effects of propofol on TREK-1 in HEK293 cells using whole cell patch clamp (**k**), and with xPLD2(**l**). **m,** Summary of TREK-1 currents showing an ∼2-fold increase when activated by propofol (25-50 µM) (n = 6) at +40 mV (± s.e.m.).

We also tested the injectable general anesthetics propofol (50 µM) (*4*). Propofol robustly activated PLD2 in N2A cells (Fig. 5d, h). If our mechanism is correct, then propofol should lead to TREK-1 activation. As predicted, propofol, robustly increased TREK-1 currents (Fig 5k) in whole cell patch-clamp. Propofol’s effect was significant (Fig. 5m, p=0.017, two tailed Student’s t test,) and co-transfection of xPLD2 with TREK-1 completely blocked the propofol specific current (Fig. 5l). Hence, PLD2 activity predicts channel function and this result suggests propofol works through the same pathway as inhaled anesthetics to activate TREK-1 albeit with less potency in C2C12 (Supplementary Fig. S6d,f).

### *In vivo* PLD dependent anesthetic sensitivity

If anesthetics disrupt membranes to activate PLD *in vivo*, then blocking PLD could block or attenuate anesthesia in an animal. Establishing anesthetic sensitivity of PLD *in vivo* would also establish the membrane and PA signaling as important up stream mediators of anesthetic action independent of a channel. To monitor sedation *in vivo*, we recorded single-animal measurements (activity and position) of *Drosophilae melanogaster* (fruit fly) in a vertically mounted chamber (Fig. 6a, Petersen unpublished data). Flies without functional PLD (PLD^null^) and wildtype (with PLD) were subjected to chloroform vapor and monitored for sedation. Sedation was determined by 5 minutes of continuous inactivity with a vertical position at the bottom of the fly chamber (Fig. 6a).

**Figure 6.**
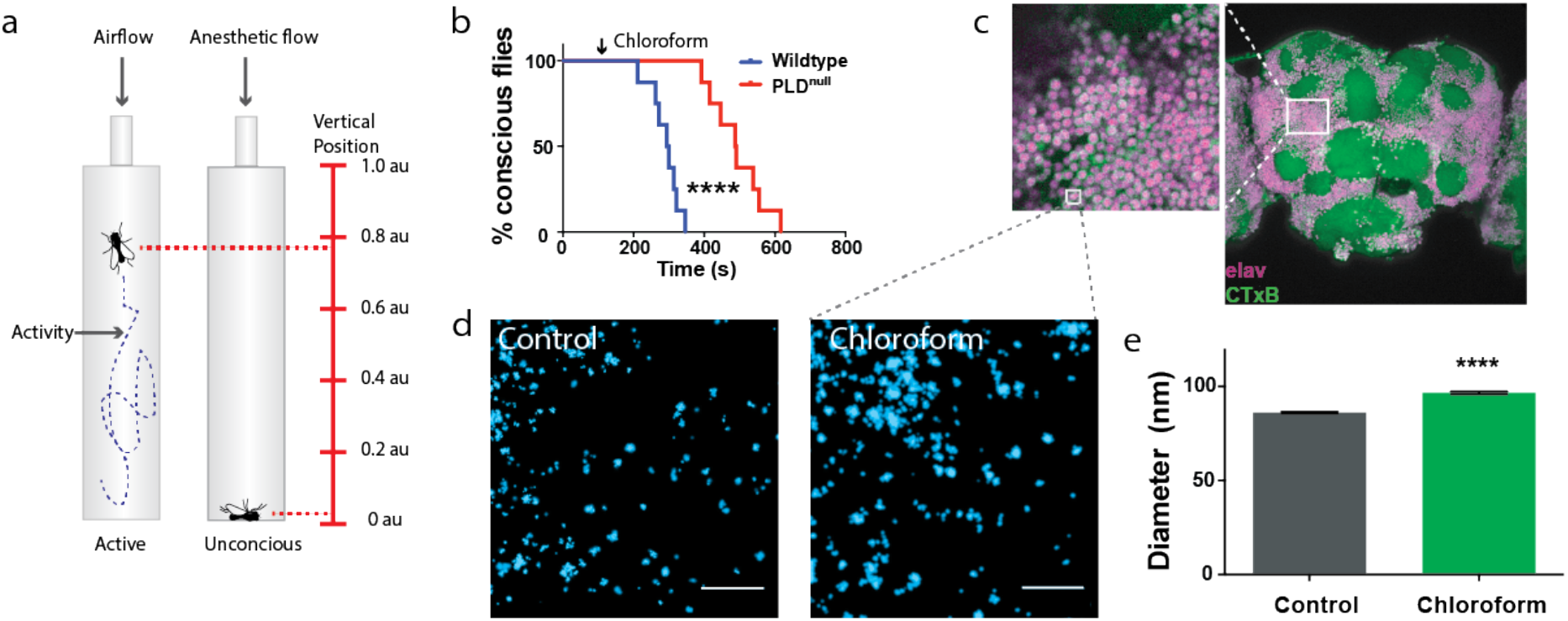
Phospholipase D (PLD) regulates anesthetic sensitivity in *Drosophila Melanogaster.* **a**, Diagram depicting the setup of the anesthetic treatment and the positional recordings of the flies. Fly positions were used to confirm anesthesia. **b**, Sedation curves showing the percent of unconscious flies over time after chloroform treatment (2.8 mmol/L of air volume) in both WT and PLD^null^ flies. p<0.00001. **c**, Confocal images showing robust labeling of GM1 domains (CtxB, green) in the membranes of labeled neurons (pan-neuronal Elav antibody, purple) of the whole fly brain. **d-e**. Flies were treated with chloroform and the GM1 domains of whole brain tissue were assayed for anesthetic induced disruption by super resolution imaging (dSTORM). **d**, Super resolution images with (right) and without (left) sedating chloroform. **e**, Quantitation of raft size diameter from fixed whole fly brain with and without chloroform. Similar to raft disruption in tissue culture, the GM1 domains expand in brains of flies treated with chloroform. Grey dotted lines indicate a hypothetical zoom compared to the low-resolution imaging of confocal in **c**.

Sedation of PLD^null^ flies with 2.8 mM chloroform required almost twice the exposure as wildtype flies (∼600 vs. 350 sec, p<0.0001) indicating a highly significant resistance to anesthesia in PLD^null^ (Fig. 6b). Almost all flies were anesthetized before the first PLD^null^ fly passed out. The concentration of chloroform is that of the vapor not the concentration in the animal. All flies eventually lost consciousness suggesting PLD sets the threshold rather than dictates anesthesia.

### Disruption of GM1 domains in whole brains

To confirm the presence of GM1 domains in brain tissue, we dissected whole brains from adult flies and labeled them with CTxB and an antibody against the pan-neuronal marker Elav. Confocal imaging (Figure 6c) of CTxB-labeled lipids (GM1, green) showed robust expression of GM1 lipids throughout the fly brain. High concentrations were observed on the membrane of the cell bodies (Figure 6c, zoom) as expected for GM1. GM1 was also observed in clusters, confirming that lipid domains also exist in the CNS of flies, making them similar in organization to our cell culture.

Lastly, we asked if anesthetic disruption of lipid rafts observed in cell culture also exists in the whole brain of an anesthetized animal. To test this, we anesthetized adult flies with chloroform, dissected their brains and characterized GM1 domains in those brains compared with a no-anesthesia control using dSTORM super resolution imaging. Consistent with cell culture, GM1 domains were expanded in fly brains treated with anesthetic (Fig. 6e). The number of domains was found to decrease by ∼10%.

## DISCUSSION

We conclude that the membrane is a target of inhaled anesthetics and that PA and disruption of palmitate mediated lipid localization contributes to anesthesia *in vivo*. Our proposed model for TREK-1 in mammalian cells is consistent with most known properties of inhaled anesthetics (Supplementary Fig. S5a) despite it utilizing an indirect mechanism through PLD2 and lipid binding sites (*11, 12, 28*). The disruption of palmitoylate mediated localization from a lipid nanodomain nicely explains how the C-terminus renders a channel anesthetic sensitive when the domain is highly charged, devoid of structure, and has no obvious hydrophobicity expected to bind an anesthetic (Supplementary Fig. S5b).

TREK-1 is not conserved in flies precluding direct downstream testing in the whole animal. These experiments will need to be done in a mammalian model. Nonetheless, our model does not rely on a single channel, rather we expect the lipid to contribute to central regulation of excitability through direct binding to multiple ion channels. Recently PA emerged as class of lipid-regulated ion channel modulating excitability and pain (*23, 34*). PA’s importance is directly supported by the finding that a single protein that modulates PA production, PLD, dramatically shifted the anesthetic threshold in an animal (Fig. 6b). The conclusion is also indirectly supported by the observation of GM1 domains throughout the brain of flies (Fig. 6c), the observed disruption of GM1 domains in the whole brain tissue of chloroform treated flies (Fig. 6e), and the modular transfer of anesthetic sensitivity to TRAAK by localization of a PA producing enzyme (Fig. 3b).

The importance of PA is evident by the fact that it is upstream of TREK-1. The latency (time delay) of PLD mixing in these experiments were dictated by our relatively slow application of anesthetic by gravity flow see methods). However theoretical estimates of latency based on distance of PLD from its substrate suggest a PLD dependent latency of ∼650 µs (*10*), much faster than our application of anesthetic ((>10 sec). In rafts that are mechanically disrupted, we have measured a PLD2 latency of TREK-1 activation with an upper limit of 2.1 ms (Petersen et. al unpublished data); we expect disruption by anesthetic is similar. PA can also affect membrane curvature and hydrophobic mismatch, and anesthetics can in theory affect these properties to regulate TREK-1(*35*), but these potential mechanism, if they contribute, appear to be insignificant compared to anesthetic activation of PLD (Fig. 2a-b).

Over the past decade anesthetic research has focused on direct binding of inhaled anesthetics to allosteric sites on proteins (*2, 4, 36*). This was not the case for TREK-1, but our results do not preclude direct binding for other channels. In some instances, the direct binding may be competition of anesthetics with lipid regulatory sites (*23*) which would be a hybrid protein/lipid mechanism. Contributing to a protein based narrative, anesthetics are also known to be enantiomer selective (*37–39*). Interestingly ordered lipids are also chiral and the ability of an anesthetic to partition into ordered domains will need to be considered in light of the mechanism presented here.

Many channels are directly palmitoylated (*22*) and they could be modulated by a change in ion channel localization. For example, the anesthetic channel γ-aminobutyric acid A receptor (GABA_A_R) gamma subunit is palmitoylated (*22*) and the alpha subunit was recently shown to bind PIP_2_ (*40*). Hence GABA_A_R is comprised of precisely the same features that render PLD2 anesthetics sensitivity. If and how these feature function in GABA_A_R is not known. Many important signaling molecules are palmitoylated including tyrosine kinases, GTPases, CD4/8, and almost all G-protein alpha subunits (*41*). Anesthetic disruption of palmitate mediated localization of these proteins likely contributes to modulation channels in a membrane dependent manner. The overall contribution of the membrane will need to be studies for each channel and regulator independently.

Lastly, we considered the biophysical effect of anesthetics on the bulk membranes or ‘non-raft’ membranes. We saw very little effect of clinical concentration of anesthetics on TREK-1 reconstituted into (DOPC) liposomes in our flux assay (Supplementary Fig. 3a-b), a mimic of bulk lipids. This result agrees with previous studies that showed the effect of anesthetics on bulk lipids is insufficient to activate a channel (*42*) at clinical concentrations despite the fact that anesthetics fluidize and thin membranes (*43*). TREK-1 is very sensitive to membrane thickness (*44*). It’s possible we failed to test an optimal thickness that is responsive in artificial systems, however, the fact that xPLD2 blocked all detectible anesthetic currents in whole cells suggests, in a biological membrane, domain disruption and TREK-1 and PLD2 translocation are the primary mechanism for anesthetic activation of TREK-1, not thinning of bulk lipids.

Our data show a pathway where anionic lipids are central mediators of anesthetic action on ion channels and these results suggest lipid regulatory molecules and lipid binding sites in channels may be effective targets for treating nervous system disorders and understanding the thresholds that govern intrinsic nerve cell excitability. Thus, the system we describe here obviously did not evolve to interact with inhaled anesthetics and a search as to what the endogenous analogue that activates this physiological system is warranted.

## ACKNOWLEDGEMENTS

We thank Andrew S. Hansen for assisting with experimental design and discussion and comments on the manuscript, Manasa Gudheti of Vutara for help with dSTORM data processing, Michael Frohman for mPLD2 cDNA, and Guillaume Sandoz for chimeric TRAAK cDNAs, and Stuart Forman for helpful discussion. This work was supported by a Director’s New Innovator Award (1DP2NS087943-01 to S.B.H.) from the NIH, a graduate fellowship from the Joseph B. Scheller & Rita P. Scheller Charitable Foundation to E.N.P. We are grateful to the Iris and Junming Le Foundation for funds to purchase a super-resolution microscope, making this study possible.

## CONTRIBUTIONS

MAP, RAL, and SBH designed the experiments and wrote the manuscript. MAP performed all electrophysiology, dSTORM, and PLD2 enzymes assays with help from ENP for dSTORM imaging, and PLD2 assays.

## SUPPLEMENTARY INFORMATION

**Supplementary Figure 1.**
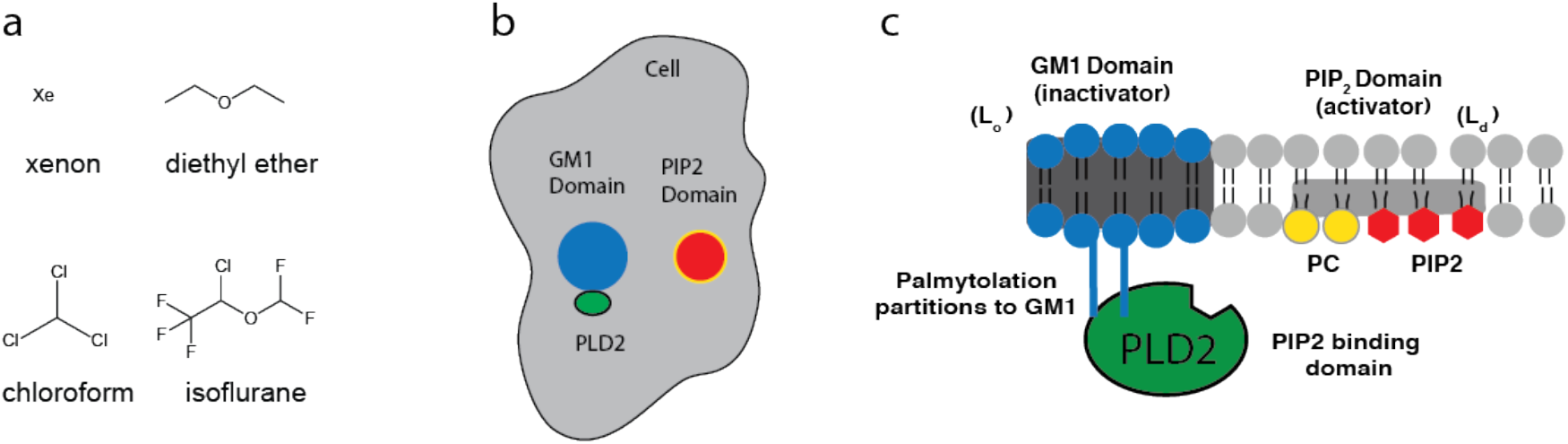
GM1 domains and PLD2 activation by substrate presentation. **a**, Chemical structures of select inhaled anesthetics are shown. Diversity ranges from xenon, a single hydrophobic atom, to chloroform, chlorinated carbon, and ether compounds. **b**, the organization of PLD2 and lipid rafts on the surface of a cell are shown (not to scale). GM-1 domains are comprised of GM1 saturated lipids and cholesterol, palmitoylation drives PLD2 into GM1 domains (blue circle) where it is sequestered away from its substrate polyunsaturated PC and PIP_2_ domains (Red circle with yellow outline). **c**, Side view of the membrane in (b) with PLD2 localized to the inner leaflet through two palmitoylation sites (blue lines). GM1 domains also commonly referred to as the liquid ordered phase (Lo) are thicker than the liquid disordered phase (Ld). We previously showed mechanical force disrupts lipid rafts and causing PLD2 translocates to the PIP_2_ domains. PLD2 has a PIP_2_ binding site and resides in equilibrium between GM1 domains and PIP_2_ domains. PIP_2_ is polyunsaturated and preferentially localized with unsaturated PC—PLD2 generates unsaturated PA and PG products through substrate presentation(*10*).

**Supplementary Figure 2.**
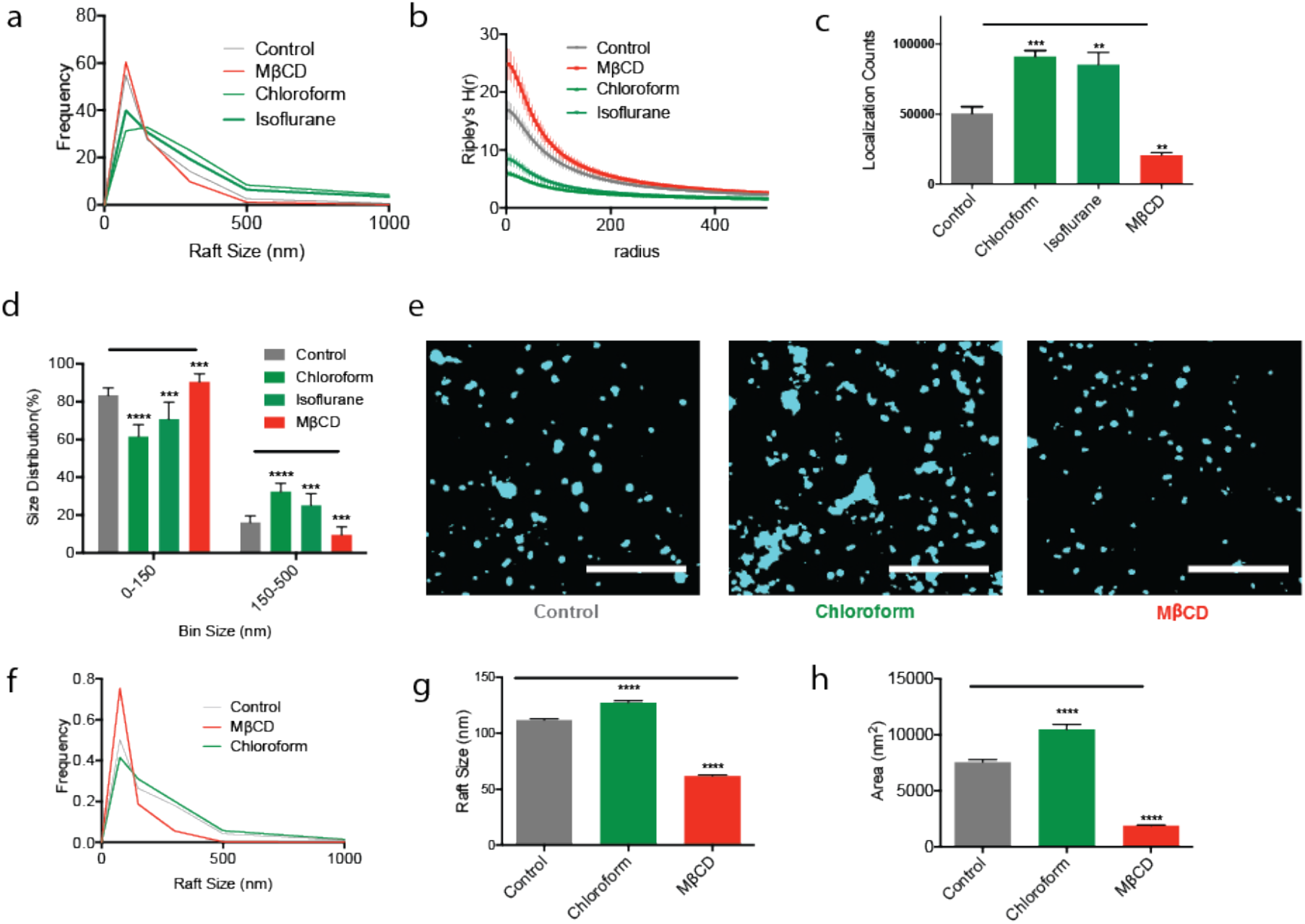
Inhaled anesthetics disrupt GM1 domains. **a**, Particle size distribution curve of the lipid rafts in N2A cells after chloroform, isoflurane, and MβCD treatments (n=10). The number of rafts smaller than 150 nm decreased and the rafts larger than 150 nm increased (see panel (d) for quantification). **b**, Derivatives of Ripley’s K-Function (H(r)) demonstrating the separation of GM1 domains with or without treatment of the inhaled anesthetics (1 mM) or MβCD (100 µM) (± s.e.m., n = 10). **c**, Bar graphs showing the amount of particle localizations acquired within the fixed frames numbers (10000) (± s.e.m., n = 5). **d**, Histograms showing the size distribution of lipid rafts from N2A cells in (a) binned from 0-150 nm and 150-500 nm. Chloroform (1mM) and isoflurane (1mM) both shift from small to large and MβCD (100 µM) shifts from large to small (± s.e.m., n = 10). Both observations are a form of disruption. **e,** Representative super-resolution (dSTORM) images of lipid rafts from C2C12 cells before and after the treatment of the inhaled anesthetics or MβCD (100 µM) (Scale bars: 1 µm). **f**, Particle size distribution curve of the lipid rafts after the chloroform, isoflurane, and MβCD treatments from multiple C2C12 cells (n = 6-8). **g-h**, Bar graphs comparing the average lipid raft sizes (**g**) and areas (**h**) quantified by cluster analysis showing the disruption of lipid raft by chloroform and MβCD (± s.e.m., n = 2844-6525). Student’s t-test results: ***P<0.001; ****P<0.0001.

**Supplementary Figure 3.**
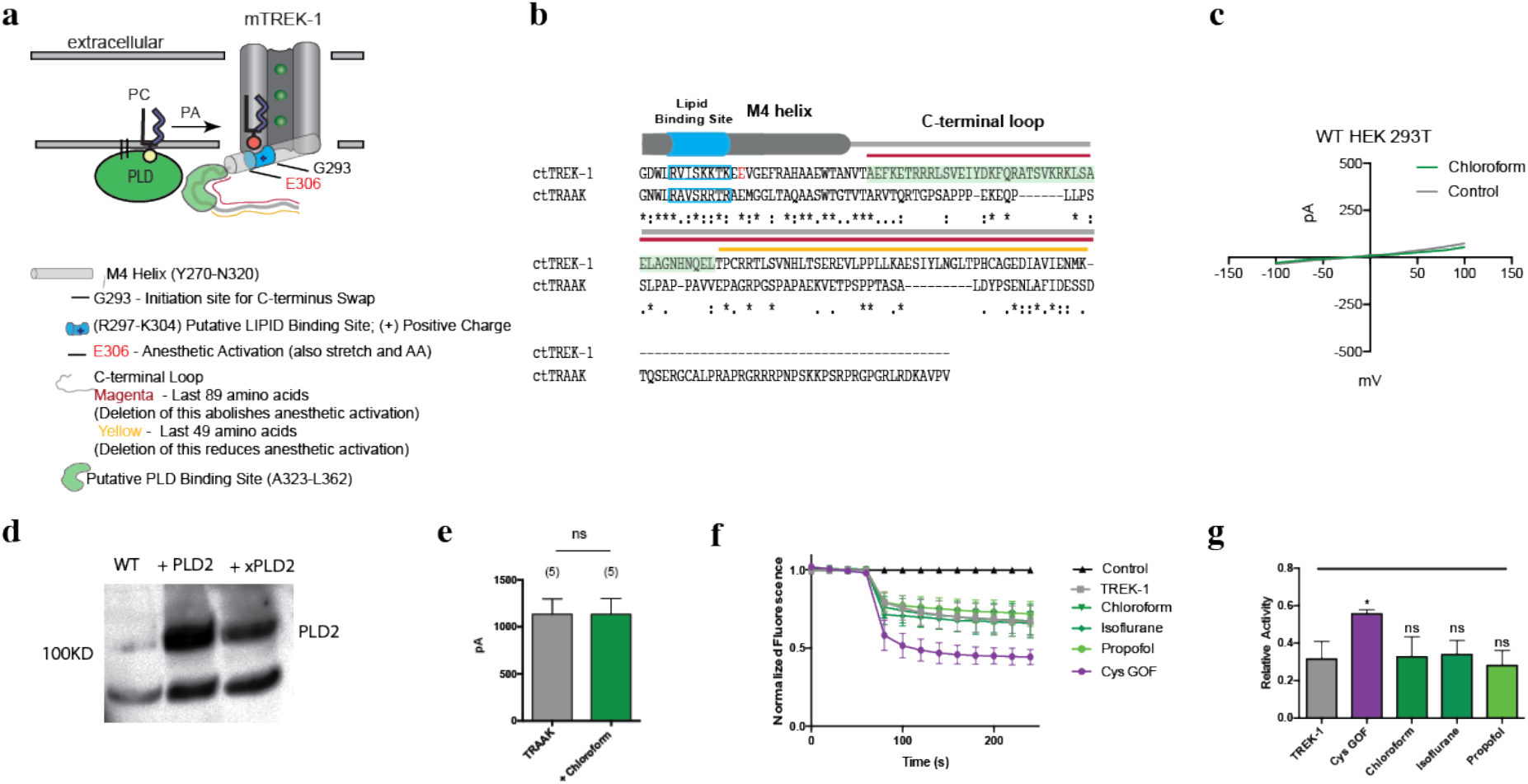
General anesthetics activate TREK-1 through PLD2. **a**, Cartoon depicting the amino acids that are known to play a role in anesthetic activation and the PLD2 dependent lipid-gating of TREK-1 channels. **b**, multiple-sequence alignment between the swapped C-terminus of mouse TREK-1 (293–411) and TRAAK (255–398) generated with Clustal Omega (EMBL-EBI). The colors and features correspond with panel (a). **c,** Current-voltage relationship (I-V) for TREK-1 in wild type HEK 293T. No change in whole cell currents by 1 mM of chloroform. **d**, Western blot showing the over expression of wtPLD2, mPLD2 and xPLD2 from transfected HEK 293T cells. **e,** Bar graph summarizing wt. TRAAK channel currents when activated by chloroform (1mM) at +40 mV (± s.e.m., n = 5) confirming earlier studies that TRAAK is anesthetic insensitive. **f,** Ion flux assay of purified TREK-1 reconstituted into DOPC (16:1) liposomes with 15 mol% DOPG anionic lipid. Anesthetics chloroform (1 mM), isoflurane (1 mM), and propofol (50 µM) had no significant effect on channel currents compare to the double cysteine gain of function mutation (Cys GOF) (± s.e.m., n = 3-5). **g,** Bar graph comparing the relative activity from **f**.

**Supplementary Figure 4.**
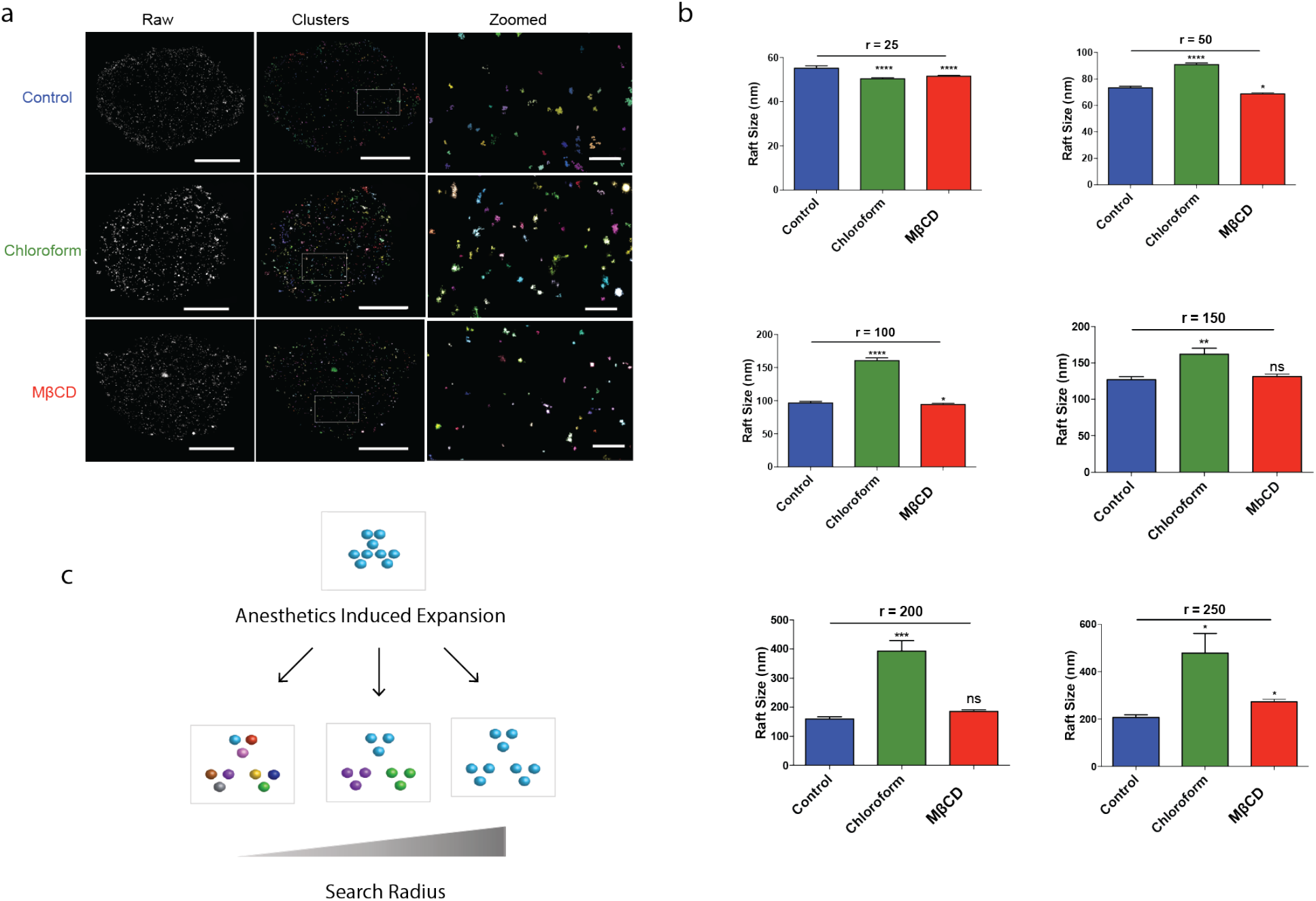
Effect of search radius on observed raft size. **a**, Representative images showing the raw images with particle localizations (left panel, scale bar = 5 μm), clustered particles (DBSCAN) to determine raft sizes (middle panel, scale bar = 5 μm), and the zoomed in (right panel, scale bar = 0.5 μm) using 100 nm search radius. Arbitrary colors are assigned to particles of the same cluster. **b**, Bar graphs showing the changes in raft sizes after anesthetic treatment at various search radius (ε) when at least 10 localizations were clustered. **c**, Cartoon depicting the size variation at different search radius. A radius that is too small will treat each particle as an individual cluster and give cluster that are too small. A search radius that is too large clusters separate molecules giving an artificially large size. The search radius of 100 nm shown in (a) and used for this study shows appropriate clustering and a significant increase in size.

**Supplementary Figure 5.**
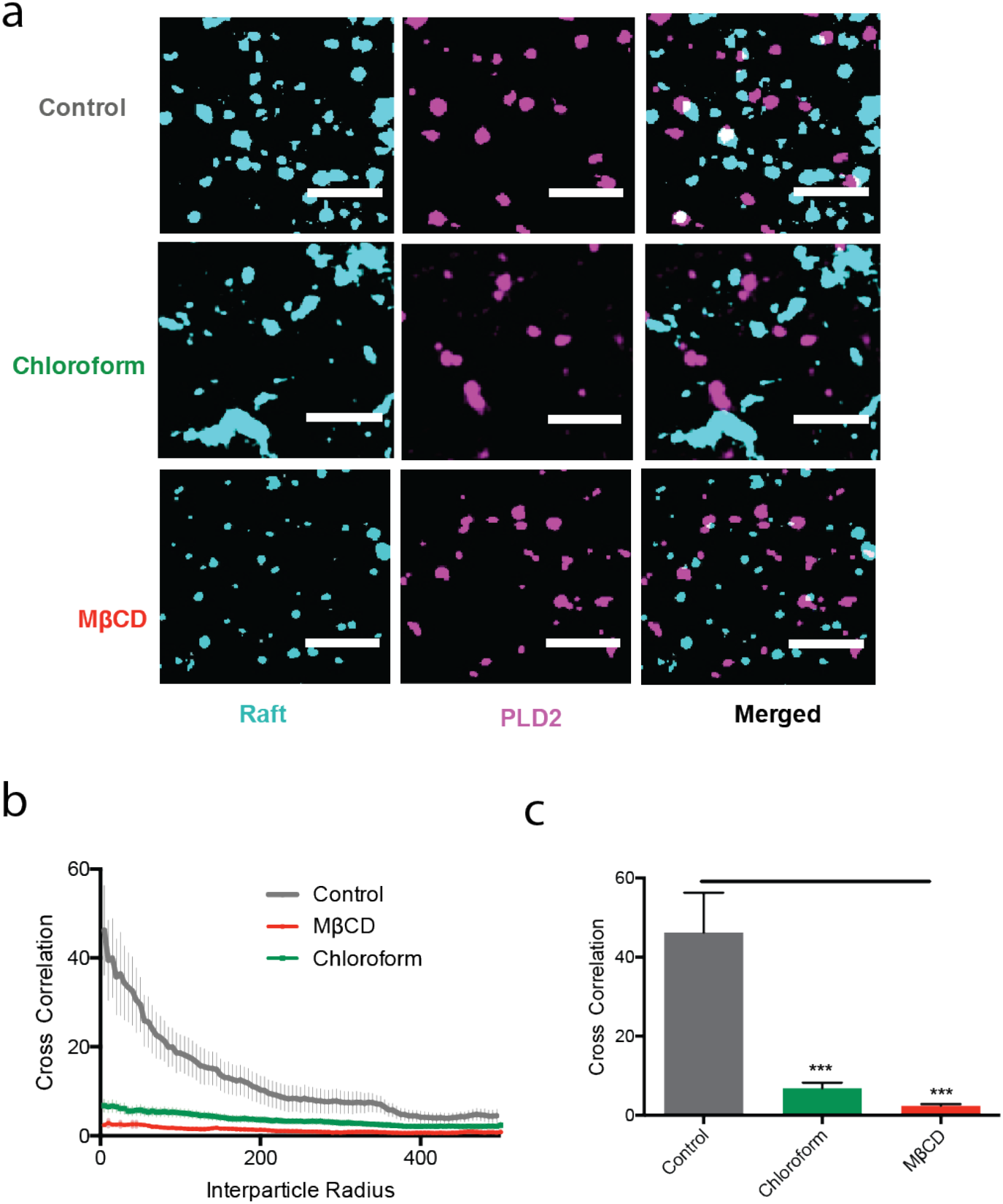
Inhaled anesthetics displace PLD2 from GM1 domains in C2C12 cells. **a**, Representative super-resolution (dSTORM) images of fluorescently labeled lipid raft (CTxB) and PLD2 before and after the treatment with chloroform (1 mM), and MβCD (100 µM) in C2C12 cells (Scale bars: 0.5 µm). **b-c**, Average Cross-correlation functions (C(r)) (**b**) and the function normalized at C(r=5) (**c**) shows chloroform (1 mM) and MβCD (100 µM), disrupt PLD2 localization into lipid raft in C2C12 cells (± s.e.m., n = 5). Student’s t-test results: *P < 0.05; ***P<0.001; ****P<0.0001; NS ≥ P.0.05.

**Supplementary Figure 6.**
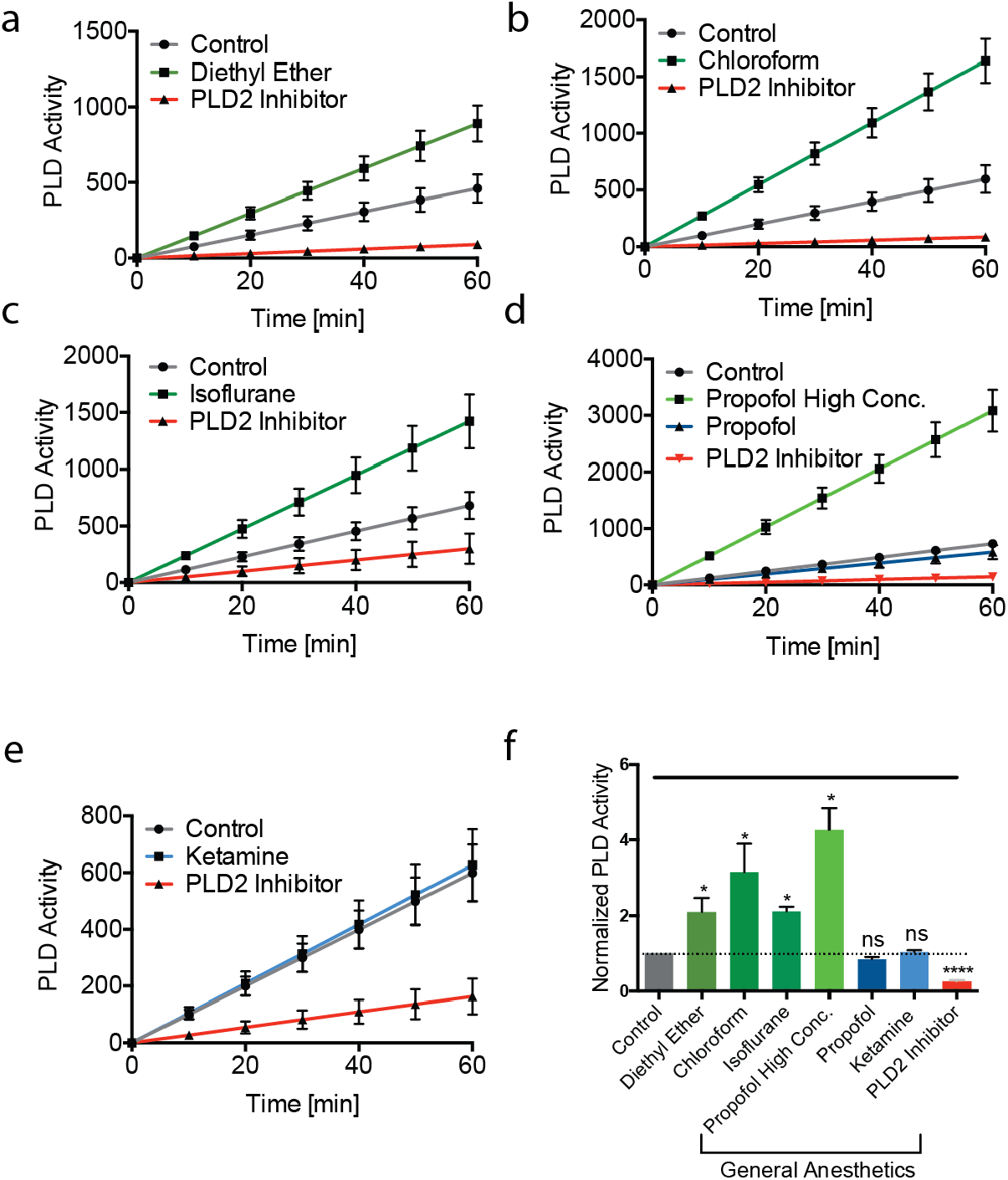
Activation of PLD2 by general anesthetics in C2C12 cells. **a-e**, Diethyl ether (1 mM) (**a**), chloroform (1 mM) (**b**), isoflurane (1 mM) (**c**), and higher concentration of propofol (400 µM), but not 50 µM propofol (**d**) (green lines) activate PLD2 compared to control cells. Ketamine (50 µM, blue line) had no observed effect on PLD2 activity. (**e**) PLD2 inhibitor (2.5-5 µM) inhibited the activity (± s.e.m., n = 4). **f**, Normalized summary effect of anesthetics on PLD2 activity at 60 min from the above experiments (± s.e.m., n = 4). Student’s t-test results: *P < 0.05; ***P<0.001; ****P<0.0001; NS ≥ P.0.05.

## Methods

### Sample preparation for Super Resolution Microscopy (d-STORM)

Super Resolution Microscopy was performed on N2A and C2C12 cells. Confluent cells were first differentiated overnight with serum-free DMEM in 8-well chamber slides (Nunc Lab-Tek Chamber Slide System, Thermo Scientific). Cells were then washed and treated with anesthetics or other drugs for 10 min. Chambers containing volatile anesthetic were tightly sealed with aluminum stickers. Cells were then chemically fixed with 3% paraformaldehyde and 0.1% glutaraldehyde in PBS for 10 min at room temperature with shaking, and the fixing solution was quenched by incubating with 0.1% NaBH_4_ for 7 min followed by three times 10 min wash with PBS. Anesthetics or the drugs were also added into the fixing solution to ensure its effect on the cell. Fixed cells were then permeabilized with 0.2% Triton-X 100 in PBS for 15 min except the cells receiving the CTxB treatment. Cells were blocked using a standard blocking buffer (10% BSA, 0.05% Triton in PBS) for 90 min at room temperature. For labelling, anti-PLD2 antibody (Cell Signaling) with 1:500 dilution and CTxB (Life Technologies) with 1:1000 dilution in the blocking buffer were simultaneously added to cells and incubated for 60 min at room temperature. Cells were then extensively washed with 1% BSA, 0.05% Triton in PBS for five times 15 min each before labeling with the secondary antibody diluted into the blocking buffer and incubating for 30 min. Prior to labeling, the secondary antibody was conjugated to either Alexa 647 (to detect CTxB raft) or Cy3B (to detect PLD2). The incubation with secondary antibody was followed by above extensive wash and a single 5 min wash only with PBS. Labeled cells were then post-fixed with the previous fixing solution for 10 min without shaking followed by three times 5 min washes with PBS and two 3 min washes with deionized distilled water. To elucidate the lipid raft disruption by anesthetics or other drugs, compounds were applied to the reaction buffer at these concentrations: chloroform (1 mM) (Fisher Scientific); isoflurane (1 mM) (Sigma); mβCD (100 μM) (Fisher); diethyl ether(1mM) (Sigma); ketamine (50 μM) (Cayman Chemicals); xenon (Praxair).

### d-STORM Image Acquisition and Analysis

Imaging was performed with A Zeiss Elyra PS1 microscope using TIRF mode equipped with an oil-immersion 63X objective. Andor iXon 897 EMCCD camera was used along with the Zen 10D software for image acquisition and processing. The TIRF mode in the dSTORM imaging provided low background high-resolution images of the cell membrane. A total of 10,000 frames with an exposure time of 18 ms were collected for each acquisition. Excitation of the Alexa Fluor 647 dye was achieved using 642 nm lasers and Cy3B was achieved using 561 nm lasers. Cells were imaged in a photo-switching buffer suitable for dSTORM: 1% betamercaptoethanol, 0.4 mg glucose oxidase and 23.8 µg Catalase (oxygen scavengers), 50 mM Tris, 10 mM NaCl, and 10% glucose at pH 8.0. Sample drift during the acquisition was corrected for by an autocorrelative algorithm(*45*) or tracking several immobile, 100 nm gold fiducial markers or TetraSpec beads using the Zen 10D software. The data were filtered to eliminate molecules with localization precisions >50 nm.

Super-resolved Images were constructed using the default modules in the Zen Software. Each detected event was fitted to a 2D Gaussian distribution to determine the center of each point spread function (PSF) plus the localization precision. The Zen software also has many rendering options including the options to remove the localization errors, outliers in brightness and size. The super-resolved images have an arbitrary resolution of 128 pixel/μm. To determine the raft size determination and the cross-correlations, the obtained localization coordinates were converted to be compatible to Vutara SRX software (version 5.21.13) by an Excel macro. Cross-correlation and raft size estimation were calculated through cluster analysis using the default analysis package in the Vutara SRX software(*10, 46–48*). Cross-correlation function *c*(*r*) estimates the spatial scales of co-clustering of two signals ─ the probability of localization of a probe to distance *r* from another probe(*49*). Raft sizes are the size of clusters determined by measuring the area of the clusters comprising of more than 10 observations.

### In Vivo PLD2 activity measurements

A nonradioactive method was performed to measure in vivo PLD2 activity as described previously (ref) (Fig. S2). Briefly, N2A or C2C12 cells were seeded into 96-well flat culture plates with transparent-bottom to reach confluency (∼ 5 x 104 per well). Then the confluent cells were differentiated with serum-free DMEM for a day and washed with 200 μL of phosphate buffer saline (PBS). The PLD assay reactions were promptly begun by adding 100 μL of working solution with or without anesthetics. The working solution contained 50 μM Amplex red, 1 U per ml horseradish peroxidase, 0.1 U per ml choline oxidase, and 30 μM dioctanoyl phosphatidylcholine (C8-PC). Anesthetics were directly dissolved into the working buffer from freshly made stocks and incubated overnight before assay reagents were added. In case of volatile anesthetics, 96-well plates were tightly sealed with aluminum sticky films after adding the reaction buffer. The PLD activity and the background (lacking cells) was determined in triplicate for each sample by measuring fluorescence activity with a fluorescence microplate reader (Tecan Infinite 200 PRO, reading from bottom) for 2 hours at 37°C with at excitation wavelength of 530 nm and an emission wavelength of 585 nm. Subsequently, PLD activity was normalized by subtracting the background and to the control activity. Data were then graphed (Mean ± SEM) and statistically analyzed (student t-test) with GraphPad Prism software (v6.0f).

### Electrophysiology

Whole cell patch clamp recordings of TREK1 currents were made from TREK-1-transfected HEK 293T cells as described previously (lab ref, Comoglio et. al. 2014). Briefly, HEK 293T were cultured in growth media [DMEM, 10% heat-inactivated fetal bovine serum, 1% penicillin/ streptomycin] in a humidified incubator (95% air and 5% CO_2)_ at 37°C. When the HEK 293T cells were ∼90% confluent, they were seed at 50% confluency per 35-mm dish containing 15mm glass coverslips coated with poly-D-lysine (1 mg/ml) to ensure good cell adhesion. The cells were then transiently transfected using X-tremeGENE (Sigma) with a total of 1μg of DNA per dish. For co-transfection of TREK1, TRAAK with PLD2 or PLD2-K758R cells were transfected with a ratio of 1:3. Human TREK1 pCEH and mouse PLD2 was kindly provided by Dr. Stephen B. Long, Sloan Kettering Institute, NY. TRAAK/Ct-TREK1 (starting at Gly 293) pIRES2eGFP and TREK1/Ct-TRAAK (starting at Gly 255) pIRES2eGFP was kindly provided by Dr. Sandoz Guillaume, iBV CNRS, Université de Nice Sophia Antipolis, France. HEK 293T cells were obtained from ATCC (Manassas, VA). Human TRAAK was a gift from Dr. Steve Brohawn, University of California, Berkeley. Transfected cells were then visualized and selected for electrophysiology 24-48 hours post transfection using green fluorescent protein. Standard whole-cell currents were recorded at room temperature with Axopatch 200B amplifier and Digidata 1440A (Molecular Devices) and measured with Clampex 10.3(Molecular Devices) at sample rate of 10 kHz and filtered at 2 kHz. The recording micropipettes were made from the Borosilicate glass electrode pipettes (B150-86-10, Sutter Instrument) by pulling with the Flaming/Brown micropipette puller (Model P-1000, Sutter instrument). The micropipette resistances were ranged from 3-7 MΩ and filled with the internal solution (in mM): 140 KCl, 3 MgCl_2_, 5 EGTA, 10 HEPES, 10 TEA pH 7.4 (adjusted with KOH). The external bath solution contained (in mM): 145 NaCl, 4 KCl, 2 CaCl_2_, 1 MgCl_2_, 10 HEPES, 10 TEA pH 7.4 (adjusted with NaOH). After the voltage offset was adjusted to zero current between the patch electrode and the bath solution, the whole cell configuration was achieved by repetitive gentle suctions on cells sealed at 1-10 GΩ. In the whole cell configuration, cells were held at −60 mV and currents were elicited by voltage steps command (at −100 to +100 mV from V_hold_ = - 60 mV) and voltage ramp commands (−100 mV to +50 mV in 5 ms). Volatile anesthetic, chloroform, was applied using a gravity-driven (5 ml/min) gas-tight perfusion systems (Valves and tubing were made of PTFE). HEK 293T cells were perfused with control solution or the test solution that contained the volatile anesthetic. Chloroform was dissolved based on the anesthetic saturation experiments that it has 66.6 mM solubility in water at 37°C(*11*). Subsequently, data were replayed and analyzed using Clampfit 10 (Molecular Devices) to generate current-voltage relationship (I-V Curve) from voltage steps protocol. Student’s *t*-test was applied to assess statistical significance using Prism6 (GraphPad software) and judged significant at *p* < 0.001. The values represented in the graphs are Mean ± SEM.

### Chanel Purification and Flux Assay

TREK-1 channel protein purification and Flux assay were done as previously described(*29, 30*). Briefly, Pichia yeast was used to express zebrafish TREK-1 (1-322 amino acids) containing GFP at C-terminus. Followed by cryo milling, the extraction of the proteins was done in dodecyl-β-d-maltoside (DDM) with protease inhibitors. The proteins were then purified on a cobalt affinity column to homogeneity followed by size exclusion chromatography (SEC). The final SEC buffer contained 20 mM Tris (pH 8.0), 150 mM KCl, 1 mM EDTA, and 2 mM DDM. All proteins were collected with a predominant monodispersed peak corresponding to the expected molecular weight (MW) of the assembled channel protein plus GFPs. This Purified TREK-1 was used to generate Proteoliposomes by mixing 1:100 TREK-1/lipids. The ratio of the Lipids (85% DOPC and 15% DOPG) were mixed, dried, and solubilized in rehydration buffer (150 mM KCl, 20 mM HEPES [pH 7.4]) and calibrated with 3 mM DDM before the channel mixing. DDM was then removed by BioBeads (Bio-Rad) and the proteoliposomes (5 μL) were sonicated and added to 195 μL of flux assay buffer (150 mM NaCl, 20 mM HEPES [pH 7.4], 2 μM 9-amino-6-chloro-2-methoxyacridine [ACMA]) in a 96-well plate at room temperature. Flux was initiated by the addition of the protonophore carbonyl cyanide m-chlorophenyl hydrazone (CCCP) (3.2 μM).

### SDS PAGE and Western Blot

HEK 293T cells were transiently transfected using X-treme gene 9 (Sigma Aldrich). After 48 hours of the transfection, cells were homogenized in the ice cold lysis buffer comprising 1.5% DDM (Anatrace), 10% Glycerol (Fisher Scientific), 1 mM EDTA (Sigma), and 1: 100 protease inhibitors (calbiochem) in 1X PBS, pH 7.5. Lysates were centrifuged at 17,000g for an hour at 4°C to collect the supernatant. Protein sample were diluted 1:1 with 2X SDS Laemmli buffer (Biorad) and incubated for 30 mins at 37°C. The denatured proteins were then separated on 4-20% SDS polyacrylamide gel (Mini-PROTEAN® TGX™, Biorad) and blotted onto polyvinylidene difluoride membranes, and probed with rabbit monoclonal antibody against PLD2 (E1Y9G, Cell Signaling). Blot signals were visualized with a AlphaImager™ Gel Imaging System (Alpha Innotech).

### Drosophila

#### General protocols

Flies were maintained in stocks at 25C. For all experiments, male flies were isolated from stocks 1-4 days old and allowed to recover in vials of no more than 10 flies for 24-48 hours before use in protocols.

#### Anesthesia in Drosophila melanogaster

Anesthesia in flies were performed by volatiles and aerosols administered with positional recording (VAAPR, Petersen unpublished data) in a custom-built narrow-width vertical chambers placed in front of a camera to monitor the fly’s positions with or without the chloroform treatment. Wildtype and PLD^null^ flies were gently loaded into the designated chamber using mouth aspiration and the hoses used for compound delivery were attached to the chamber. Flies were allowed to habituate in the chamber while preparing treatment mixtures (∼15 min). Mixtures without chloroform contains only water with methylcyclohexanol (MCH) (1: 250), an aversive odor to increase baseline activity of the flies. Chloroform were delivered at known concentrations by calculating the partial pressures of the anesthetic in known dilutions of solvent. Both air control and chloroform air were passed through flow meters to control the overall flow rate of 290 mL/min/chamber. Air was given to all flies for 2 min to record baseline activity/position after which the chloroform was applied to the experimental flies. Data was analyzed by the Noah.py program for position and activity(ref). Time of half maximal anesthesia (T50i) values were obtained using a best-fit variable slope curve; sedation curves were analyzed using the Mantel-Cox test. Curve fitting and statistical tests were determined using GraphPad Prism 6.

#### Brain Imaging

Flies were placed into control vial or vial containing a chloroform-soaked tissue. Flies were removed and brains were isolated and placed into buffer containing PBS or PBS+chloroform on ice. After dissections, brains were transferred to a buffer containing PBS with 3% paraformaldehyde and 0.1% glutaraldehyde and allowed to fix while rocking overnight at 4C. The following day the tissue was rinsed with PBS containing 0.5% TritonX-100 and 0.5% BSA. This buffer was used for all additional steps unless otherwise stated. Tissue was rinsed 2X for 1hr after which buffer with additional 30mg of BSA was added and allowed to rock at rt for 1.5 hours. Buffer was removed and identical buffer containing primary antibodies (DSHB, Rat-Elav-7E8A10, 1:200) was added to the tissue after which it was rocked for 3 hr at rt and then overnight at 4C. The following day buffer was used to rinse the tissue as was done previously. Secondary antibody (Jackson ImmunoResearch, 112-606-072, 1:500) was then applied in a similar manner as before but rocking at 4C was extended to 5 days. On the 5^th^ day, the tissue was rinsed once again 2X and then in PBS alone. Tissue was then prepared for imaging. CTxB (1:500) was treated as a secondary antibody. Super-res imaging was performed as described above. For whole brain imaging confocal imaging was used after mounting the tissue onto a coverslip using standard protocols.

